# ATAD2 BRD mediates liquid-liquid phase separation of ATAD2 to promote histone acetylation

**DOI:** 10.64898/2026.03.08.708675

**Authors:** Chang Shu, Zhou Gong, Yuanfang Wang, Yusen Zhang, Maili Liu, Xu Zhang, Danyun Zeng

**Affiliations:** Key Laboratory of Magnetic Resonance in Biological Systems, State Key Laboratory of Magnetic Resonance and Atomic and Molecular Physics, National Center for Magnetic Resonance in Wuhan, Wuhan Institute of Physics and Mathematics, Innovation Academy for Precision Measurement of Science and Technology,Chinese Academy of Sciences, Wuhan,430071; University of Chinese Academy of Sciences,Beijing,100049; Wuhan National Laboratory for Optoelectronics, Huazhong, University of Science and Technology,Wuhan,430071; Optics Valley Laboratory, Wuhan,430074

## Abstract

ATAD2 possesses a C-terminal bromodomain (BRD) that plays a critical role in recognizing and binding to acetylated lysine residues. However, because the native intracellular structure of ATAD2 remains poorly defined, the mechanisms by which the ATAD2 BRD recruits acetylated histones and the regulatory pathways involved are not yet understood. In this study, we report that the ATAD2 BRD mediates the formation of liquid-liquid phase separation (LLPS) of ATAD2 in cells. This phase separation promotes the process of histone H4 acetylation, leading to the up-regulation of C-MYC, CCND3, and ATF2 gene expression and the facilitation of chromatin remodeling. Our findings elucidate a vital function of ATAD2, wherein BRD-mediated LLPS drives histone acetylation to promote cellular chromatin remodeling.

## Introduction

ATAD2 is a nuclear-localized chromatin regulator, also recognized as an oncogenic transcriptional co-activator^[1]^, that is highly expressed across various malignancies^[2]^. Structurally, ATAD2 comprises two AAA+ ATPase domains and a C-terminal Bromodomain (BRD). The ATAD2 BRD is characterized by a conserved fold of four α-helices (αZ,αA,αB,αC) and two flexible loops (ZA and BC loops). Together, these loops and specific $\alpha$-helical residues form a hydrophobic cavity, termed the RVF shelf, which serves as a docking site for the acetylated N-terminal tails of histones^[3]^.As a central “reader” of acetylated histones, ATAD2 BRD is not merely a static structural component but a core driver of cellular chromatin remodeling. Emerging evidence suggests that bromodomain-containing proteins often aggregate on chromatin through multivalent weak interactions, forming highly dynamic liquid-liquid phase separation (LLPS) condensates. This transition into a condensed physical state effectively concentrates diffuse transcription factors and chromatin-modifying enzymes at specific genomic loci, thereby dramatically enhancing the efficiency of biochemical reactions.While it is established that the two AAA+ ATPase domains of ATAD2 form a hexameric complex responsible for ATP binding and hydrolysis^[4]^, the precise structural configuration of the BRD within the full-length protein remains elusive. ATAD2 BRD plays a pivotal role in DNA replication, transcriptional regulation, and the modulation of histone modifications^[5]^, and its dysregulation is linked to diverse human cancers, including breast, liver, gastric, colorectal, and lung adenocarcinomas^[6][7]^. Consequently, it serves as a significant prognostic indicator in oncology^[8]^.As a reader of histone acetylation, the ATAD2 BRD is indispensable for chromatin remodeling^[9]^. Although ATAD2 BRD reportedly exhibits a high affinity for H4K5ac^[5]^—a modification critical for the cellular response to chromatin damage^[10]^—the precise mechanism by which ATAD2 BRD recruits acetylated histones and regulates histone acetylation patterns in vivo remains poorly understood. Here, we report that ATAD2 BRD mediates the formation of ATAD2 liquid-liquid phase-separated condensates. This phase separation facilitates the process of H4 histone acetylation and modulates downstream gene transcription, thereby driving functional chromatin remodeling.

## Results

### ATAD2 BRD Undergoes Liquid-Liquid Phase Separation (LLPS) In Vitro

ATAD2 BRD exhibits the capacity to undergo liquid-liquid phase separation (LLPS) in solution. Although previous studies have noted that ATAD2 BRD forms aggregates in solution^[11]^, the specific morphology of these assemblies has not been characterized. Using Transmission Electron Microscopy (TEM), we observed that ATAD2 BRD forms spherical assemblies at both 4-hour and 12-hour time points, with the diameter of these structures increasing progressively over time (Fig. 1a).To further characterize these spherical assemblies, we conjugated ATAD2 BRD with the red fluorescent tag 5(6)-TAMRA SE. Under confocal microscopy, the protein formed spherical, droplet-like puncta. Furthermore, Fluorescence Recovery After Photobleaching (FRAP) analysis demonstrated a rapid recovery of the ATAD2 BRD fluorescent signal within the bleached area, indicating that these spherical assemblies possess internal liquid-like fluidity (Fig. 1b, c, d). Collectively, these results demonstrate that ATAD2 BRD undergoes LLPS in solution.

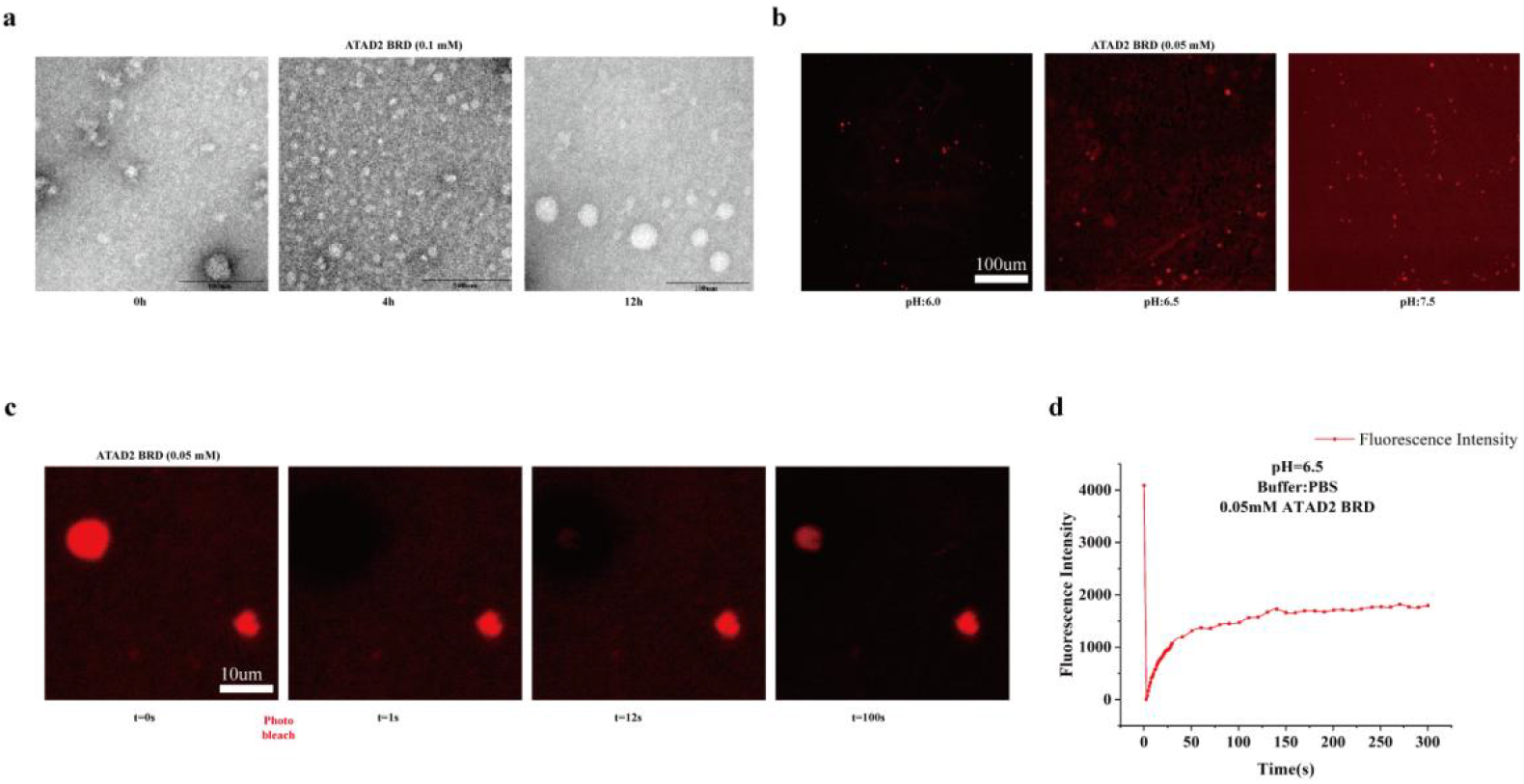

### ATAD2 Forms LLPS Mediated by Its Bromodomain

The functional execution of ATAD2 requires the synergistic action of its AAA+ ATPase domains and its Bromodomain (BRD)^[12]^. Given that the isolated BRD undergoes liquid-liquid phase separation (LLPS) in solution, we investigated whether it dictates the phase behavior of the full-length ATAD2 protein. To evaluate the inductive capacity of ATAD2 for LLPS, we purified and fluorescently labeled recombinant full-length ATAD2 (residues 1–1390, tagged with EGFP). We observed that ATAD2 readily formed spherical droplets in vitro (Fig. 2b). Subsequently, we transfected HeLa cells with a plasmid encoding EGFP-tagged full-length ATAD2 (ATAD2-EGFP) and monitored its morphology in live cells via confocal microscopy. ATAD2-EGFP partitioned into spherical puncta within the nuclei of HeLa cells (Fig. 2c). Fluorescence Recovery After Photobleaching (FRAP) analysis confirmed the liquid-like properties of these droplets, as evidenced by the rapid recovery of the fluorescent signal post-bleaching (Fig. 2d, e).

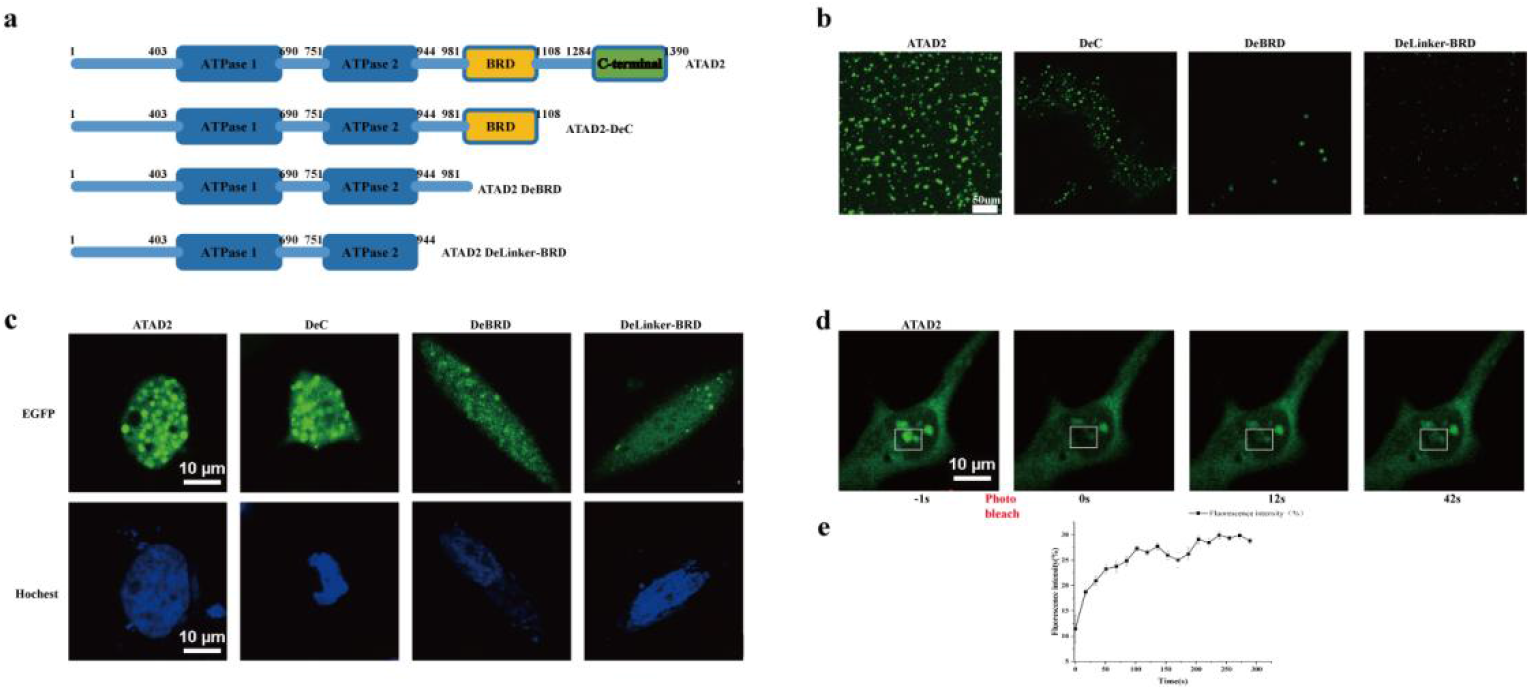

We next sought to determine whether ATAD2 LLPS is specifically mediated by the BRD. We generated and purified a series of C-terminally EGFP-tagged ATAD2 truncations: DeC (C-terminal truncation following the BRD), DeBRD (deletion of the entire BRD), and DeLinker-BRD (deletion of the BRD along with the linker connecting it to the AAA domain) (Fig. 2a). At room temperature, only the DeC construct formed robust spherical droplets. In contrast, the DeBRD mutant produced significantly fewer and smaller droplets compared to DeC; this residual phase separation may be attributed to the linker region, which contains low-complexity intrinsically disordered regions (IDRs)^[13][14]^[15]. Notably, the DeLinker-BRD mutant failed to form droplets entirely, appearing as irregular aggregates (Fig. 2b). Consistent with these in vitro findings, live-cell imaging of HeLa cells expressing these truncations showed that DeC formed prominent spherical droplets, while DeBRD showed diminished droplet formation, and DeLinker-BRD showed none (Fig. 2c). Importantly, these differences in phase behavior were not due to variations in protein expression levels.

To further validate the requirement of the BRD, we treated cells with GSK8814, a potent ATAD2 BRD inhibitor that competitively blocks the binding of acetylated histones to the BRD^[16]^. With increasing concentrations of GSK8814, ATAD2-mediated LLPS was progressively attenuated and eventually abolished. Collectively, these findings demonstrate that the formation of LLPS by ATAD2 is primarily mediated by its Bromodomain.

### LLPS of ATAD2 is accompanied by a concentration-dependent conformational transition

Our investigations into the ATAD2 bromodomain (BRD) revealed that increasing protein concentrations trigger a distinct shift in its circular dichroism (CD) spectra. Specifically, the characteristic α-helical peak at 190 nm disappears, replaced by absorption peaks indicative of β-sheet structures. These data suggest that ATAD2 BRD undergoes a structural transformation as concentration increases. This observation was further corroborated by Thioflavin T (ThT) fluorescence assays, which showed a time-dependent increase in β-sheet content until reaching an equilibrium that correlates positively with protein concentration.We further monitored the intracellular conformation of ATAD2 post-LLPS. Consistent with our in vitro findings, ATAD2 exists primarily in a β-sheet-dominant conformation within cellular droplets. While the AAA+ ATPase domain of ATAD2 is reported to function as a hexamer characterized by α-helical structures, the gradual shift of the BRD toward a β-sheet conformation suggests that the overall structural transition of ATAD2 is mediated by its BRD.We also evaluated the impact of temperature, concentration, and pH on this conformational change and discovered that the transition is sensitive to redox conditions. Specifically, inhibiting the formation of disulfide bonds within ATAD2 BRD significantly suppresses its structural transformation. Liquid-state NMR (^1^H-^15^N HSQC) spectroscopy showed that at a concentration of 0.1 mM, the characteristic peaks of ATAD2 BRD decayed rapidly. However, the addition of 15 mM DTT to create a reducing environment dramatically slowed the decay of these HSQC signals. This indicates that preventing disulfide bond formation inhibits the conformational transition of the BRD.In conclusion, these results demonstrate that ATAD2 LLPS is mediated by its BRD and progresses alongside a conformational shift regulated by disulfide bond formation.

### ATAD2 BRD facilitates histone acetylation through liquid-liquid phase separation (LLPS)

The ATAD2 bromodomain (BRD) comprises a conserved bundle of four α-helices and two loop regions. Within this structure, the ZA loop and portions of the α-helices form a hydrophobic pocket, known as the RVF shelf, which specifically recognizes acetylated lysine (Kac) residues on histone tails, exhibiting the highest binding affinity for H4K5ac.To investigate whether the droplets formed by ATAD2 BRD influence its binding to acetylated histones and the subsequent acetylation process, we utilized H4K5 as a model substrate (peptide sequence: SGRGKGGKGLGKGGAKRHRK) in an acetylation assay mediated by the acetyltransferase EP300. By supplementing the reaction with varying concentrations of ATAD2 BRD (Fig. 3a), we monitored the reaction kinetics. After one hour of incubation, the production of Coenzyme A (CoA) in groups containing ATAD2 BRD was significantly higher than in the control group (0 mM ATAD2 BRD), demonstrating a positive correlation between the degree of H4K5 acetylation and ATAD2 BRD concentration (Fig. 3b).Simultaneously, fluorescence microscopy revealed that LLPS occurred within the system upon the addition of ATAD2 BRD (Fig. 3c), suggesting that the formation of these droplets promotes histone acetylation. To verify that this enhancement is directly driven by LLPS, we employed SDA (Sulfo-NHS-diazirine) to photo-crosslink ATAD2 BRD, thereby inhibiting its ability to undergo phase separation (Fig. 3c). When the crosslinked ATAD2 BRD was added to the H4K5 acetylation system, the acetylation levels remained comparable to the control group (without ATAD2 BRD). This further confirms that ATAD2 BRD facilitates histone acetylation specifically through the formation of LLPS.In a cellular context, we performed a component analysis of ATAD2-formed droplets. By utilizing cell lysis followed by sucrose gradient centrifugation, we isolated ATAD2 droplets and confirmed the enrichment of H4K5ac within these condensates via immunoblotting (Fig. 3e).

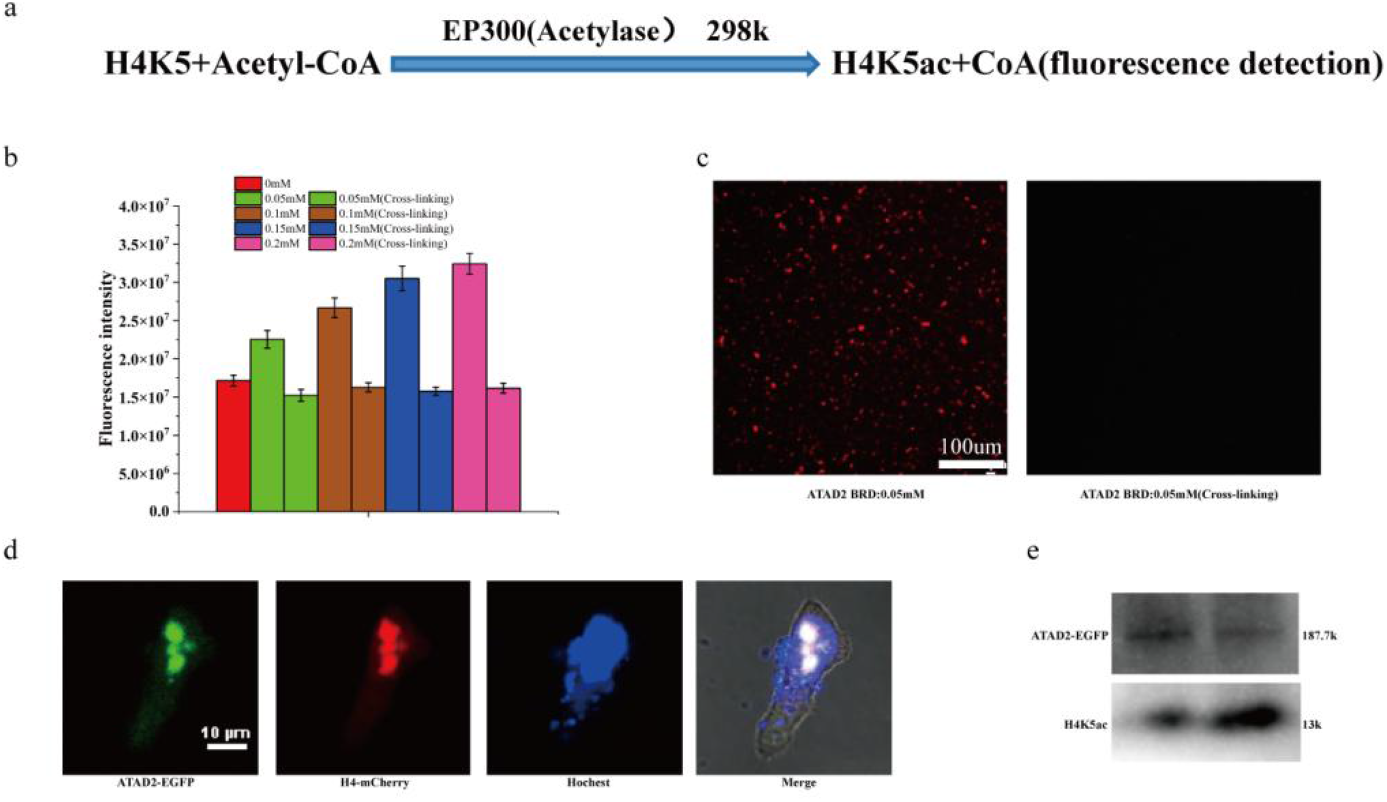

### BRD-mediated ATAD2 LLPS facilitates the chromatin remodeling process

When ATAD2 exists within multimers or high-molecular-weight complexes, the inter-domain interactions within ATAD2 enhance the binding efficiency to histone H4 acetylation sites. To determine the impact of intracellular ATAD2-mediated LLPS on H4 acetylation, we evaluated H4 acetylation levels across three groups: ATAD2 (HeLa cells exhibiting LLPS), DeBRD (HeLa cells lacking LLPS), and NC (normal HeLa cells). We observed that H4K5ac levels in the ATAD2 group were significantly higher than those in the NC group (Fig. 4a, b), validating that the formation of ATAD2 LLPS promotes the histone H4 acetylation process. Interestingly, while H4K5ac levels in the DeBRD group were lower than in the ATAD2 group, they remained higher than the NC group. This suggests that the AAA+ ATPase domain may facilitate ATP hydrolysis to maintain H4K5ac levels above baseline, yet the absence of LLPS prevents reaching the maximal acetylation observed in the ATAD2 group.

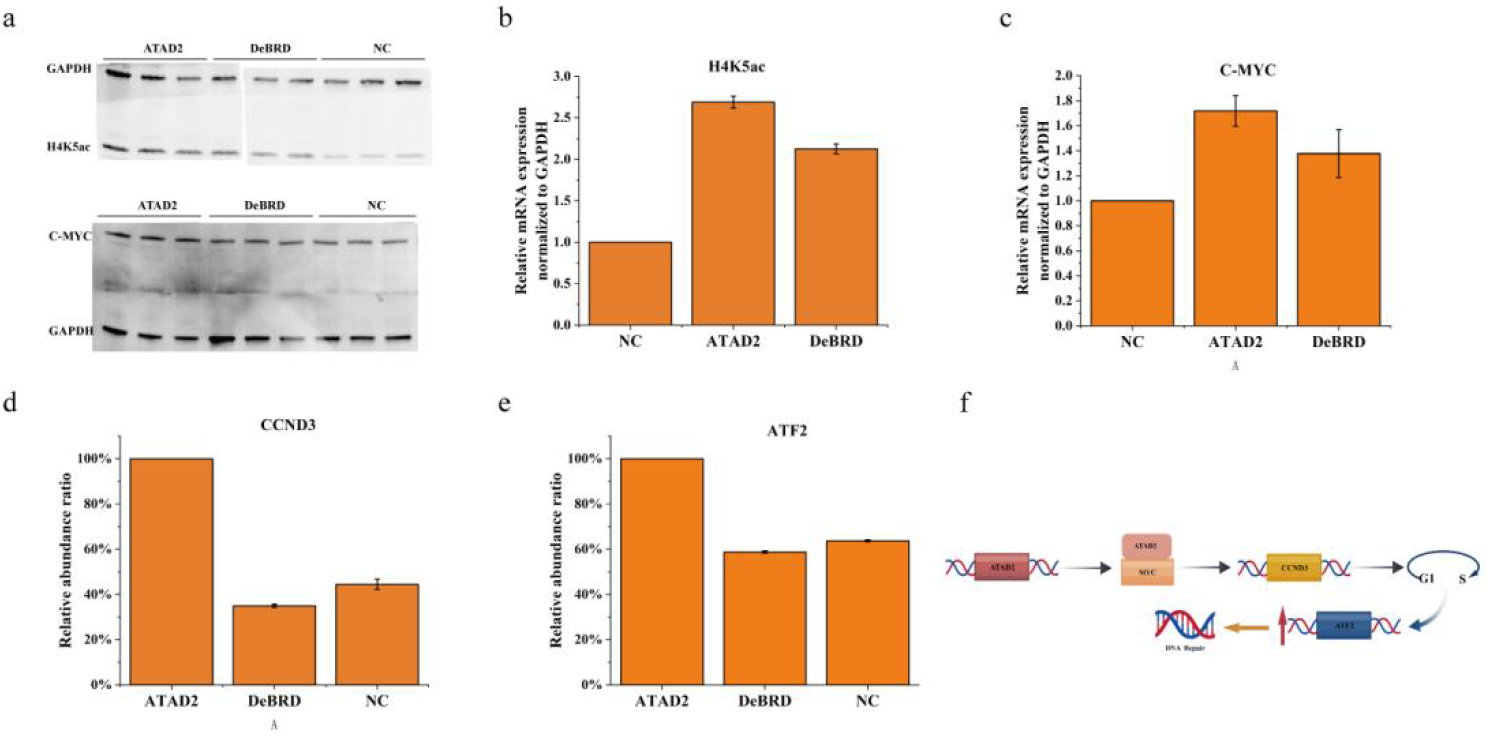

Furthermore, we observed that the transcriptional levels of C-MYC, CCND3, and ATF2 were upregulated as a result of ATAD2 LLPS (Fig. 4c, d, e). Mechanistically, ATAD2 indirectly promotes CCND3 expression by activating oncogenes such as C-MYC, thereby accelerating the cell cycle transition from G1 to S phase^[20][21]^[22]. This indicates that ATAD2 LLPS heightens the activation of C-MYC and CCND3 expression, facilitating the G1-S transition and subsequent chromatin remodeling. We also detected an upregulation in the transcription of ATF2, a transcription factor involved in the DNA damage response that is highly expressed during the G1-S phase^[23][24]^ (Fig. 4f), further corroborating the acceleratory effect of ATAD2 LLPS on the cell cycle.

In summary, these results support the conclusion that BRD-mediated ATAD2 LLPS promotes the expression of C-MYC and CCND3, driving the G1-to-S phase transition and accelerating chromatin remodeling.

## Discussion

The structural understanding of ATAD2 has long been confined to isolated studies of its AAA+ ATPase and bromodomain (BRD). While recent structural reports of full-length ATAD2 reveal that the AAA+ ATPase domain functions as a hexamer, the inherent high flexibility of the BRD has hindered the determination of its authentic conformation within the full-length protein. Our study demonstrates that full-length ATAD2 undergoes liquid-liquid phase separation (LLPS), a process mediated by its BRD. Experiments utilizing various truncation mutants further validate the indispensable role of the BRD in driving ATAD2 phase separation.

The induction of ATAD2 LLPS correlates with a significant increase in histone H4 acetylation and the subsequent activation of downstream gene transcription. Specifically, the upregulation of C-MYC indirectly promotes CCND3 expression, thereby facilitating the G1-to-S phase transition and promoting chromatin remodeling. Interestingly, we also observed that ATAD2 LLPS enhances the expression of ATF2. Since ATF2 binding is known to inhibit acetyltransferase activity^[25]^, this suggests a potential negative feedback mechanism. In this model, ATAD2 LLPS triggers a rapid surge in cellular acetylation levels, which in turn upregulates ATF2 to suppress acetyltransferase activity, thereby maintaining homeostatic acetylation balance. The precise regulatory nuances of this feedback loop warrant further investigation.

In conclusion, our research proposes a novel mechanism by which ATAD2 regulates intracellular acetylation through phase separation. These findings offer fresh perspectives on how ATAD2 participates in chromatin remodeling and provide a new functional context for understanding the authentic intracellular conformation of ATAD2.

## Materials and Methods

### Protein Expression and Purification of Recombinant ATAD2 BRD

The recombinant plasmid pET21a(+)-ATAD2 BRD was transformed into R(s) competent cells. Following transformation, cells were resuscitated in antibiotic-free LB medium at 37℃ and 220 rpm for 30 min, harvested by centrifugation, and plated onto LB agar containing kanamycin. After overnight incubation at 37℃, single colonies were inoculated into 1000 mL of kanamycin-resistant LB medium and cultured at 37℃ and 220 rpm until the OD600 reached 0.8-1.0. Protein expression was induced by the addition of 1.0 mmol/L isopropyl-β-D-1-thiogalactopyranoside (IPTG) followed by overnight incubation at 17℃ and 220 rpm.Cells were harvested and resuspended in 50 mL of lysis buffer (500 mmol/L NaCl, 50 mmol/L HEPES, 3 mmol/L imidazole, 5% glycerol, pH 7.5). After lysis via high-pressure homogenization at 1000 bar, the lysate was clarified by centrifugation at 20000 rpm for 40 min. The supernatant was filtered through a 0.22 um membrane and loaded onto a Ni-NTA affinity column. The column was equilibrated with Buffer A (500 mmol/L NaCl, 50 mmol/L HEPES, 5 mmol/L imidazole, 5% glycerol, pH 9.0). Target proteins were eluted using a linear gradient of 10%–70% Buffer B (containing 300 mmol/L imidazole).The eluted fractions were concentrated to 1-2 mg/mL, supplemented with TEV protease, and dialyzed against Buffer A overnight to remove the affinity tag. The cleavage mixture was re-applied to the Ni-NTA column equilibrated with Buffer A to remove the protease and uncleaved proteins. The purified ATAD2 BRD was finally concentrated, desalted, and stored at 4℃.

### FRAP

For liquid-liquid phase separation (LLPS) visualization, ATAD2 BRD was labeled with TAMRA-SE (5-carboxytetramethylrhodamine, succinimidyl ester; Thermo Fisher). TAMRA-SE was dissolved in DMSO (1 mM) and stored at -20℃ in the dark. ATAD2 BRD was buffer-exchanged into PBS and incubated with TAMRA-SE at a 1:5 molar ratio for 1.5 h at room temperature, protected from light. Excess dye was removed by repeated ultrafiltration (30 kDa, 3900 rpm, 30 min) until the filtrate was colorless.The labeled protein was quantified using a BCA assay and stored at 4℃. For imaging, TAMRA-labeled ATAD2 BRD was mixed with 10% PEG8000 in PBS and equilibrated on a glass slide for 3 min. Images were acquired using a Nikon confocal microscope. A 561 nm laser was used to bleach the center of the protein droplets, and fluorescence recovery was recorded at 2-second intervals for 300 seconds. Data analysis was performed using Origin 2021b. All experiments were conducted in triplicate.

### Western Blotting

HeLa cells were maintained in DMEM supplemented with 10% fetal bovine serum (FBS) and 1% penicillin-streptomycin at 37℃ in a 5% CO_2_ atmosphere. Transient transfection was performed using Lipofectamine 3000 according to the manufacturer’s protocol. After 24 h, cells were harvested and lysed via ultrasonication on ice for 5 min. The lysate was centrifuged (13000 rpm, 15 min), and the supernatant was collected.Samples (8 uL supernatant mixed with 2 uL loading buffer) were resolved by SDS-PAGE at 150V for 40 min. Proteins were transferred to NC membranes using an ACE automated transfer system. Membranes were blocked with 5% BSA for 1 h at room temperature and incubated with primary antibodies (ABclonal; 1:2000 dilution) overnight at 4℃. After three 5-minute washes with 5x TBST (diluted 1:5), the membranes were incubated with HRP-conjugated secondary antibodies (1:20,000 dilution) for 1 h at room temperature in the dark. Following four 5-minute washes with TBST, protein bands were visualized using a chemiluminescence kit (Beyotime) according to the manufacturer’s instructions.

